# PySight: plug and play photon counting for fast intravital microscopy

**DOI:** 10.1101/316125

**Authors:** Hagai Har-Gil, Lior Golgher, Shai Israel, David Kain, Ori Cheshnovsky, Moshe Parnas, Pablo Blinder

## Abstract

Imaging increasingly large neuronal populations at high rates pushed multi-photon microscopy into the photon-deprived regime. We present PySight, an add-on hardware and software solution tailored for photon-deprived imaging conditions. PySight more than triples the median amplitude of neuronal calcium transients in awake mice, and facilitates single-trial intravital voltage imaging in fruit flies. Its unique data streaming architecture allowed us to image a fruit fly’s olfactory response over 234 *×;* 600 *×;* 330*µm*^3^ at 73 volumes per second, outperforming top-tier imaging setups while retaining over 200 times lower data rates. PySight requires no electronics expertise nor custom synchronization boards, and its open-source software is extensible to any imaging method based on single-pixel (bucket) detectors. PySight offers an optimal data acquisition scheme for ever increasing imaging volumes of turbid living tissue.

## Introduction

Multi-photon laser scanning microscopy (MPLSM) provides a glimpse into the functioning mammalian brain with sub-cellular resolution (5, 24, 25). Recent improvements in optical microscope design, laser sources and fluorophores (22) have extended the use of MPLSM to challenging applications such as imaging of very large neuronal populations (21) and fast volumetric imaging (12). These applications face a common inherent limitation: a given rate of photon detections is spread across a rapidly increasing number of voxels sampled per second. In the resulting photon-deprived regime, several photodetector-induced noise sources reduce the correlation between the total electrical charge acquired from the photodetector, and the actual number of photons it has detected (6). Photon counting improves the signal to noise ratio (SNR) by thresholding electrical current fluctations into uniform photon detection events (6, 15). This improvement is particularly useful in neuronal calcium and voltage imaging, where a small increase in imaging conditions has a large impact on spike detectability (8). Implementing photon counting for multidimensional intravital microscopy has so far required custom electronics and a relatively high level of expertise. We introduce here *PySight*, an add-on solution that seamlessly embeds photon counting into most existing multi-photon imaging systems. It combines commercial, off-the-shelf hardware with open-source software, tailored for rapid planar and volumetric imaging. PySight uniquely time-stamps each photon detection event with 100 picoseconds accuracy, resulting in a modest data throughput while exceeding the spatio-temporal resolution of existing volumetric imaging setups (3, 9, 12).

## Results

### A. System architecture

The anatomy of conventional multi-photon systems involves, among others, a pulsed laser source, beam steering elements and their auxiliary optics, a collection arm with one or more photomultiplier tubes (PMTs), pre-amplifiers and an analog-to-digital acquisition board (25). Figure 1a depicts such a system with PySight photon-counting add-on. Electrical pulses following photon detections in each PMT are optionally amplified with a high-bandwidth preamplifier (TA1000B-100-50, Fast ComTec). The amplified pulses are then conveyed to an ultrafast multiscaler (MCS6A, Fast Comtec) where a software-controlled discriminator threshold determines the voltage amplitude that will be accepted as an event. The arrival time of each event is registered at a temporal resolution of 100 picoseconds, with no deadtime between events. This basic setup, along with the PySight software package, is sufficient for multi-dimensional imaging. Figure 1b shows summed time-lapses of the same field of view (FOV) at different times for digital and analog photon detection schemes, while detailed instructions on system setup and use are provided as a supplementary protocol. Converting the detected photon arrival times into multi-dimensional time series is a matter of interpolating the corresponding instantaneous position of the laser beam focal point within the sample. PySight computes the difference between photon arrival times and the respective synchronization signals from the laser beam steering elements (Supplementary Figure S1). Using a few key inputs from the user like the scanning mirror’s frequency, it then builds a multi-dimensional histogram and populates each voxel with the respective number of photons that were collected when the laser beam focused on it. The histogram can either be rendered and viewed directly, or be processed further by registering it to the moving brain’s frame of reference, and computing quantitative metrics about its content (neuronal activity, blood flow, etc.) (22). As rendering takes place off-line, experimental monitoring is done by routing one of the multiscaler outputs (`SYNC`, Figure 1a) to the analog-to-digital card of the existing system. The output of this channel is similar to that of a high-end discriminator, which already reduces PMT-dependent noise.

**Fig. 1.**
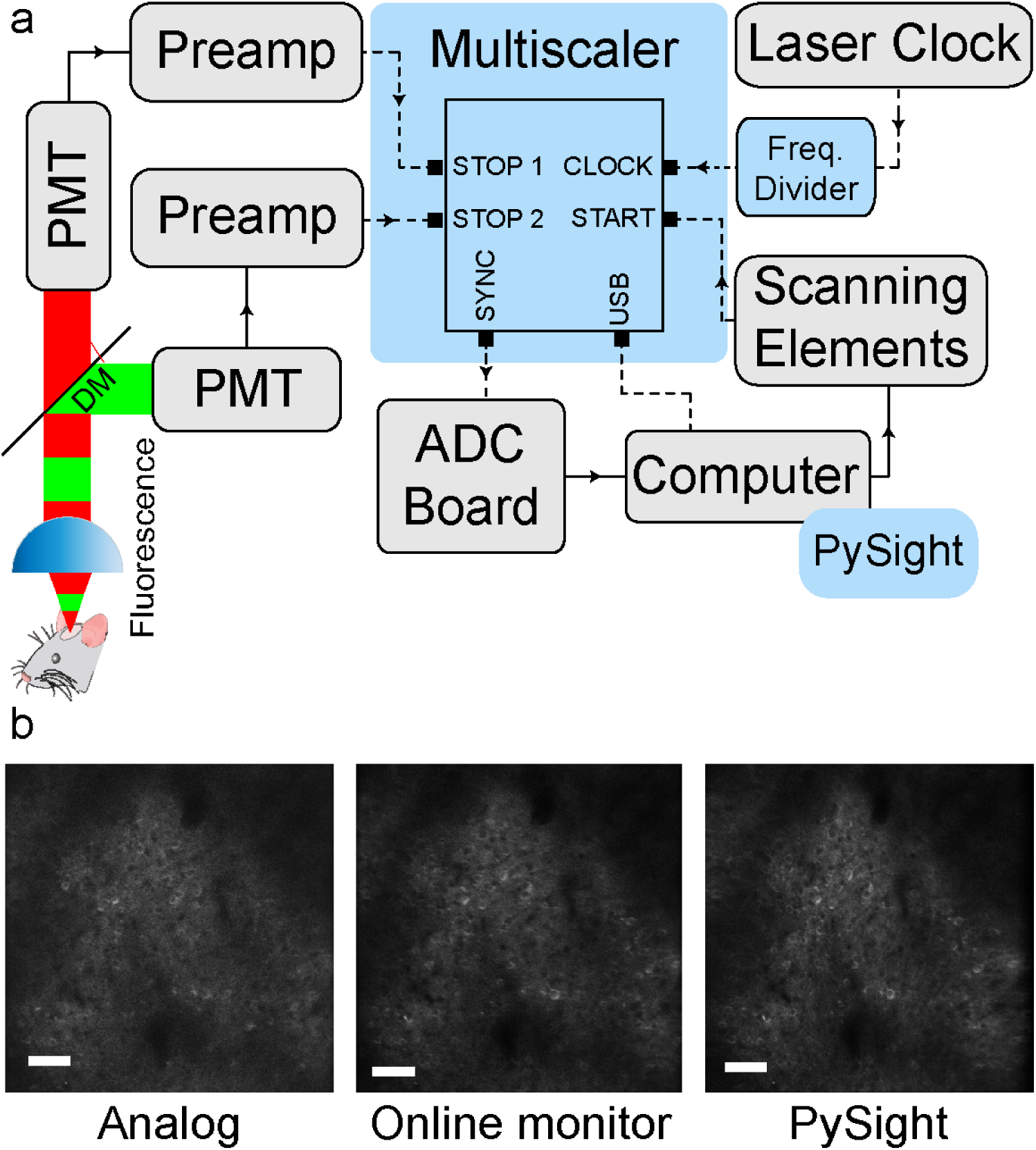
The imaging setup of the proposed system and representative *in-vivo* images taken from an awake mouse expressing a genetically encoded calcium indicator under a neuronal promoter (Thy1-GCaMP6f) a) A typical multi-photon imaging setup, depicted in gray, can be easily upgraded to encompass the multiscaler and enable photon-counting acquisition (blue). The output of the PMTs, after optional amplification by fast preamplifiers, is relayed to the multiscaler’s analog inputs (`STOP1` and `STOP2`) where it’s discretized, time-stamped and logged. Finally, the PySight software package, provided with this article, processes the logged data into multi-dimensional time series. Additionally, the multiscaler’s `SYNC` port can output the discriminated signal for a specific PMT, enabling simultaneous digital acquisition and monitoring of the discriminated signal through the analog imaging setup. b) Images produced by analog and digital acquisition schemes. Images were summed over 200 frames taken at 15 Hz. Scale bar is 50 *µ*m. DM - dichroic mirror. PMT - photomultiplier tube. Preamp - preamplifier. ADC - analog to digital converter.

### B. PySight improves calcium imaging in awake mice

We used PySight to image neurons expressing GCaMP6f under the Thy-1 promoter in awake mice, this within a normal photon flux regime, and compared its performance to analog integration within the same FOV, imaging conditions, and during the same imaging session. We analyzed both analog and PySight-generated movies (two mice, four fields of view per acquisition type) using CaImAn, a calcium analysis framework (18). PySight’s noise suppression allowed us to use about five times higher PMT gains (control voltage of 850mV vs. 650mV in analog imaging), which gave rise to improved calcium imaging and analysis even under normally encountered imaging conditions (Figure 2). Following peak detection filtering, which resulted in a mean firing rate of about 0.2 Hz for both acquisition types, we found that calcium imaging with PySight produces considerably higher Δ*F/F* calcium transients when comparing spike-like events from the entire FOV (Figure 2e); analog: median of 16% for 311 cells, PySight: median of 57% for 324 cells, *p <* 0.0001, Mann-Whitney test). Accordingly, the mean calcium transient was 247.1% high with PySight and 26.1% using analog imaging (*p <* 0.001, Welch’s two-sided t-test, same cell numbers). Additionally, PySight’s calcium transients have faster rise times than their analog counterparts (Figure 2f); 0.40 s for 1933 events vs. 0.42 s for 2112 events, *p <* 0.001, Mann-Whitney test). Having direct access to photon counts reduced the mean data throughput by a factor of 7.5-11.5 compared to the same number of 16-bit pixels during analog acquisition, and allowed us to estimate the relationship between Δ*F/F* and the number of detected photons (Supplementary Figure S2): on average a Δ*F/F* of 100% corresponds to 5.28 photons per second per *µm*^2^.

**Fig. 2.**
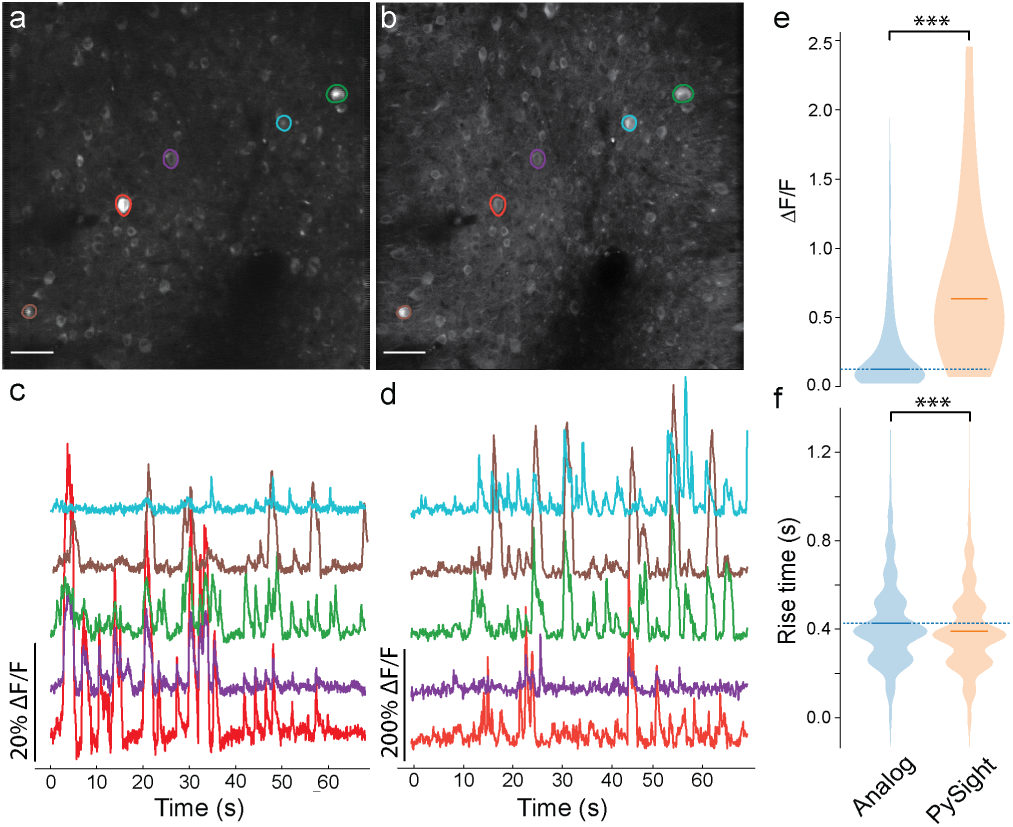
Intravital calcium imaging. (a-d) Intravital calcium imaging with analog imaging (a,c) and PySight (b,d). Notice the 10-fold larger Δ*F/F* scale for PySight-generated calcium transients (b,d). e) The distribution and median of the Δ*F/F* values for spike-like events. Median of PySight-generated calcium transients are 3.6 higher than those in analog imaging (57% for 324 cells, 16% for 311 cells, *p <* 0.0001, Mann - Whitney test). f) The distribution and median of the calculated rise times for spike-like events in the calcium traces. PySight-generated rise times are shorter than those in analog imaging (0.40 s for 1933 events vs. 0.42 s for 2112 events, *p <* 0.001, Mann - Whitney test). Scale bar for (a) and (b) is 50 *µm*.

### C. PySight enables rapid intravital volumetric imaging

Next, we utilize the exquisite temporal precision (100 picoseconds) of PySight’s hardware for ultrafast volumetric imaging. We implemented the fastest continuous axial scanning scheme available today, based on an ultrasonic variofocal lens (TAG lens), in a setup and fashion similar to Kong and coworkers (12). Figure 3 demonstrates volumetric calcium imaging of olfactory brain areas in live *Drosophila* using a TAG lens. The TAG lens modulates the effective focal depth of the excited volume sinusoidally at a rate of 189 kHz, with no synchronization to the scanning mirrors that steer the beam laterally. Despite the asynchronous scanning, PySight successfully resolves the volumetric origin of each collected photon (see Figure 3 and Supplementary Figure S3), based on synchronization TTL pulses delivered by the TAG lens driver and the planar scanning software. This simple solution obviates the phased-locked loops and photodiode-based synchronization apparati devised in earlier studies for analogous data acquisition systems (12, 17).

**Fig. 3.**
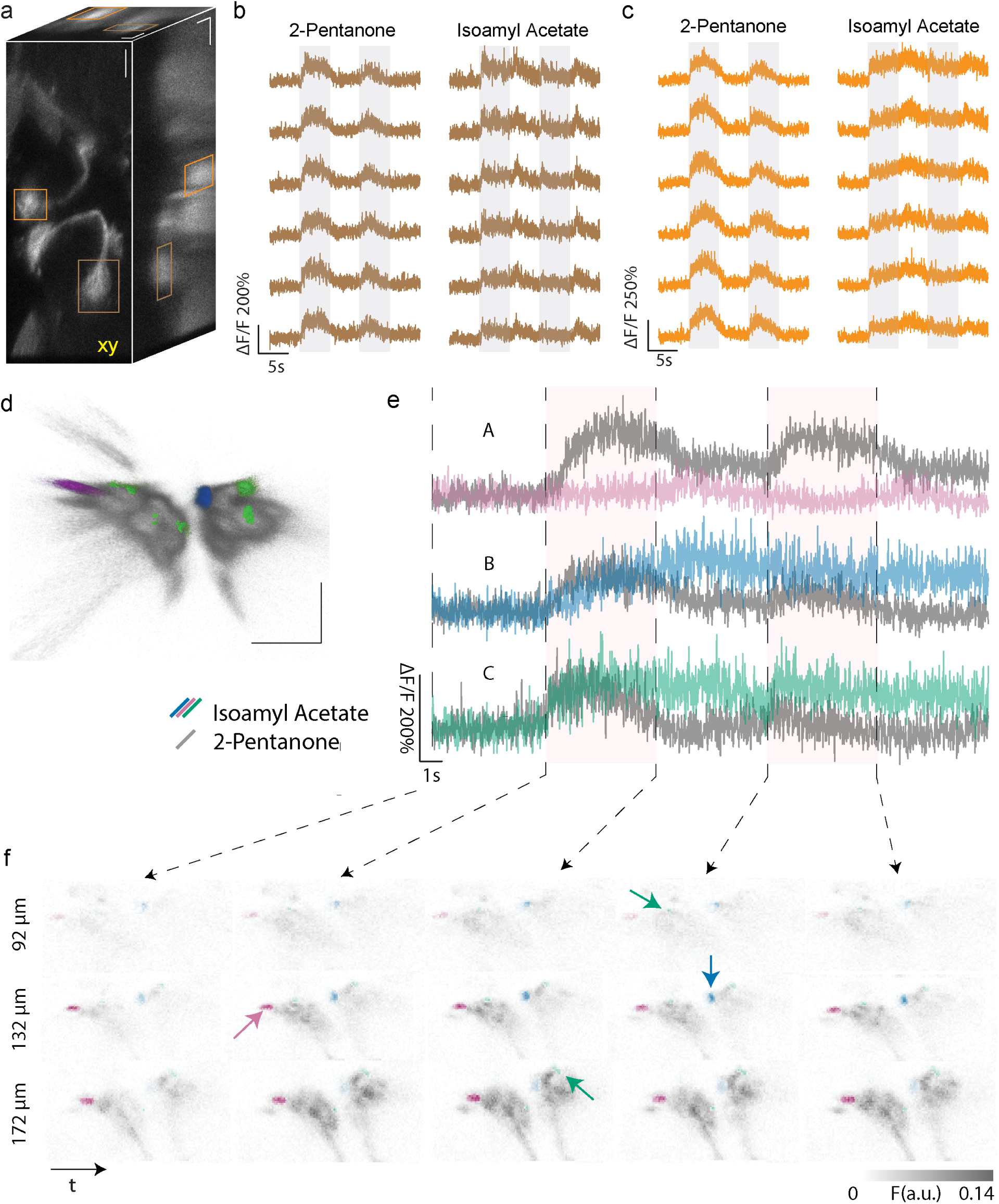
Intravital volumetric imaging of a live GCaMP6f-expressing fruit fly brain, at a rate of 73.4 volumes per second, while the fly was exposed to two different odors. Multiview projections of the imaged volume to the XY (left), XZ (top) and YZ (right) planes, summed across the 234 *×;* 600 *×;* 330*µm*^3^ volume over 33.5 seconds. b,c) Single-trial fluorescence variations over 25 seconds in the left lateral horn (b, brown) and right antennal lobe (c, orange). The fruit fly was exposed to odor puffs during seconds 5-10 and 15-20. While the lateral horn and antennal lobe responded similarly to the 2-Pentanone puffs with prolonged onset responses, their responses consistently diverged for isoamyl acetate puffs. (d-f) Glomeruli-specific odor response dynamics within the antennal lobes. d) Volume rendering of the antennal lobes with artificial coloring of glomeruli A (magenta), B (blue) and C (green). Scale bar equals 100 *µm*. (e) Volumetric GCaMP6f traces (Δ*F/F*) from distinct glomeruli in the fly’s antennal lobes were acquired at 67.2 volumes per second over a volume of 110 *×;* 257 *×;* 330*µm*^3^. The mean response to Isoamyl Acetate and 2-Pentanone odor puffs is traced in colorful and gray traces, respectively. The pink rectangles mark the duration of the odor puffs. Glomerulus A exhibits a graduated weakly-adpating response to 2-Pentanone contrasted by a weak response to isoamyl acetate, whereas glomerulus B exhibits a sustained response to isoamyl acetate, that peaks well after the odor puff has ceased. While glomeruli A and B were identified according to their morphological structure, glomeruli C were identified according to their preferential response to isoamyl acetate, which was consistently stronger than their response to 2-Pentanone. f) Montage of axial slices of the antennal lobes, highlighting the localization of the glomeruli rendered in (d) and traced in (e) within the antennal lobes. Each slice spans 6.6 *µm* in z and 5 seconds in time, axially spaced 40 *µm* apart. The glomeruli are separable laterally, axially and by their response dynamics to both odors, and are marked by colorful arrows.

We imaged the antennal lobes, mushroom bodies and lateral horns of a GCaMP6f-expressing *Drosophila*, while the fly was repeatedly exposed to two different odors: 2-Pentanone and isoamyl acetate. The imaged volume, spanning 234 *×;* 600 *×;* 330*µm*_3_, was large enough to image all six non-coplanar olfactory regions simultaneously (see Figure 3 and Supplementary Figure S3). At an imaging rate of 73.4 volumes per second, the resulting mean data throughput of 5.1 MB/s was 221 times smaller than the data throughput incurred by 8-bit analog imaging at the same voxelization (200 *×;* 512 *×;* 150 voxels in *xyz*). While the lateral horn and antennal lobe responded similarly to the 2-Pentanone puffs with prolonged onset responses, their responses consistently diverged for isoamyl acetate puffs (see Figure 3b-c and Supplementary Figure S3). Zooming in on the antennal lobes, distinct glomeruli were identified based on their morphology, response dynamics and odor preference (see Figure 3d-f). Glomerulus A exhibits a graduated weakly-adpating response to 2-Pentanone contrasted by a weak response to isoamyl acetate, whereas glomerulus B exhibits a sustained response to isoamyl acetate, that peaks well after the odor puff has ceased. While glomeruli A and B were identified according to their morphological structure, glumeruli C were identified according to their preferential response to isoamyl acetate, which was consistently stronger than their response to 2-Pentanone. Glomeruli C were identified in the anterior and medial edges of both antennal lobes (labeled in green in Figure 3d-f). All these response dynamics are absent from an empty volumetric region of interest, selected right on top of the left antennal lobe (see Supplementary Fig. S3). Hence PySight is capable of resolving distinct laminar dynamics simultaneously sampled by the variofocal lens, rather than merely extending the effective depth of field using Bessel beams, as demonstrated earlier (14).

### D. Planar intravital voltage imaging

To demonstrate the portability and ease of use of our add-on hardware, we performed voltage imaging in a different laboratory across campus using a commercial multi-photon system (DF-Scope, Sutter Inc). Detection –in single trials– of genetically encoded voltage indicators (GEVI) responses is an exceptionally challenging application due to their small fractional changes, fast photobleaching, and low replacement in the cell membrane (1). Under two-photon imaging, detection of single spikes required the use of photon counting, either through direct measurement of fluorescence changes or using fluorescence life time imaging (1). Recent advanced in the development of GEVIs allowed recording responses to visual stimuli in the fly brain (23), albeit requiring trial averaging. Here, PySight resolved single-trial odor responses, *in-vivo*, in neurites expressing ASAP2f on the fly antennal lobe (Figure 4).

**Fig. 4.**
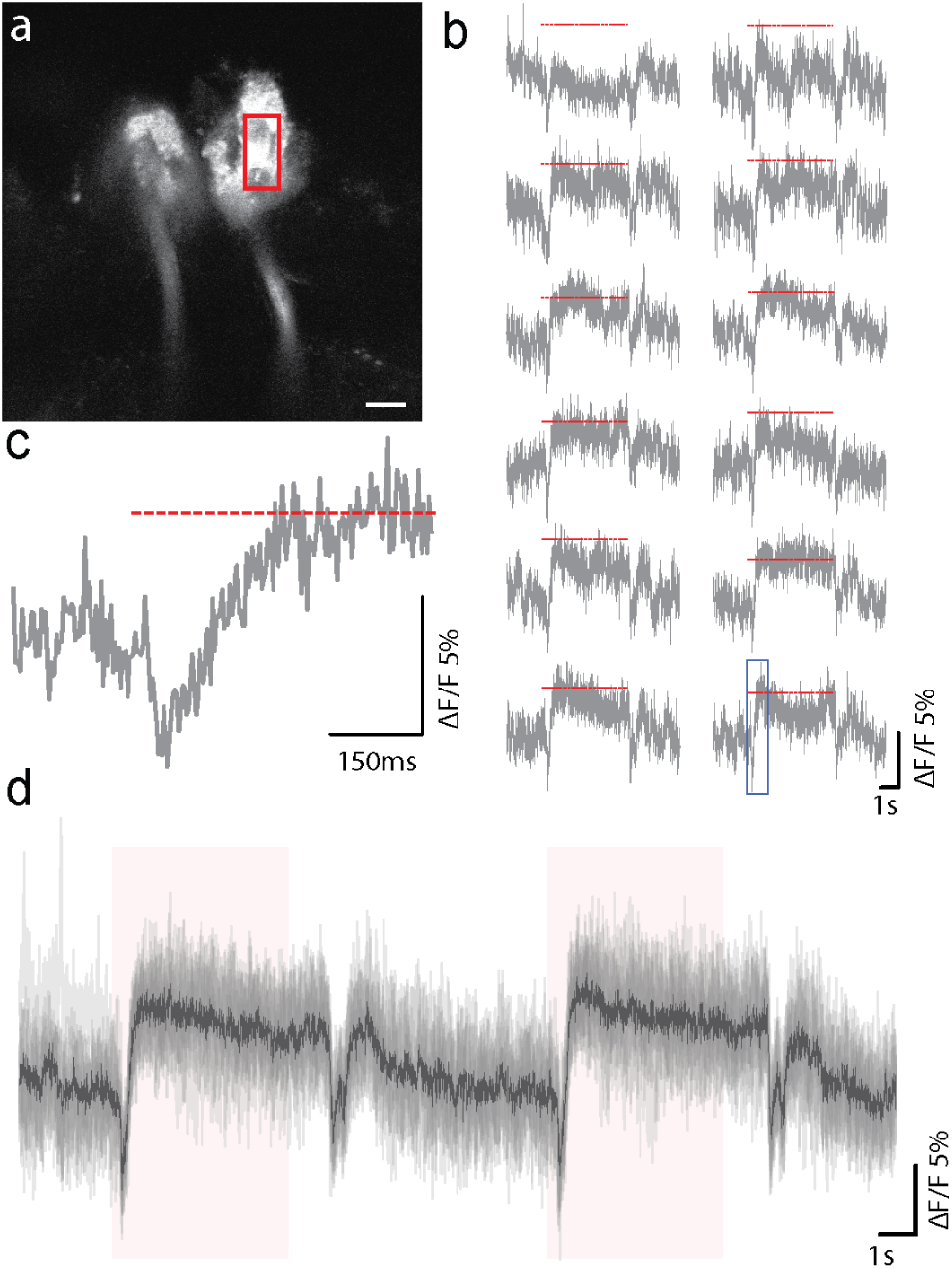
Odor responses of a GEVI-marked antennal lobe. a) Representative image of baseline GEVI signal with a ROI marked over the area used to measure responses over the antennal lobe. b) Examples of twelve different single-trial responses to a five second odor stimuli (isoamyl acetate). The red bar marks the odor puff duration. c) Zoomed-in trace, marked by the blue rectangle in (b). d) Overlaid traces (gray) with their mean (black) and the puff duration (red). Scale bar represent 50 *µm*.

## Discussion

The neuroscience community is steadily striding towards high-throughput, rapid volumetric imaging of multiple brain regions (22, 24). To achieve these feats, data acquisition hardware has to maintain single-photon sensitivity for tracking dim features, while still handling spouts of photons emitted from sparse bright features, this over lengthy imaging sessions. Analog DAQ systems lump the electrical charge sampled per voxel, thereby degrading SNR and imposing a trade off between spatio-temporal resolution and data throughput. Conversely, the data throughput with the time-stamping photon counter used here scales only with the number of detected photons, alleviating the need to sacrifice resolution in order to converge at realistic write speed or storage space. We have shown here how a suite of commercial off-the-shelf hardware and our open-source software dramatically boosts neuronal calcium imaging in awake mice and voltage imaging in the fly brain. PySight has more than tripled the median amplitude of neuronal calcium transients in calcium imaging of cortical layer 2-3 of awake mice (Figure 2), owing in part to the thresholding step that eliminates most of the variance in PMT current fluctuations in the presence and absence of detected photons. Its added value is expected to grow with imaging depth and speed. Moreover, PySight facilitated rapid continuous volumetric imaging of the fly’s olfactory system with unprecedented spatio-temporal resolution, dissecting the odor response dynamics of individual glomeruli in 4D (Figure 3d-f).

Users may also synchronize the photon counter with their laser pulses through its reference clock input (1, see methods), enabling several applications such as fluorescence life-time imaging (6, 9, 15), image restoration (7), temporal demultiplexing of interleaved beamlets (3, 9) and post-hoc gated noise reduction. Moreover, as PySight provides direct access to the photon count in each voxel, it allows implementing photon mis-detections correction algorithms that can further increase the dynamic range of the rendered images (6, 10, 16, 20). Finally, although PySight has been built around specific hardware, the open-source code can handle any list of photon arrival times through a well-documented application interface (see Supplementary Figure S1c and Supplementary Note L. It is extensible to any imaging method based on single-pixel (bucket) detectors, including fast compressive bioimaging (13, 19). Therefore, PySight’s versatility allows multiple experimenters to use the same rendering code independent of experimental setups. Together with its superior noise suppression and unmatched spatio-temporal precision, PySight offers an optimal data acquisition scheme for ever increasing imaging volumes of turbid living tissue.

## Methods and Materials

### E. Animals and Anesthesia

All imaging experiments and surgical procedures were approved by the Tel Aviv University ethics committees for animal use and welfare and followed pertinent IACUC and local guidelines. Adult male mice from the C57BL/6J-Tg(Thy1-GCaMP6f)GP5.5Dkim/J transgenic line were used. These mice constitutively express the GCaMP6f calcium indicator, mainly in L2/3 and L5 neurons(2). Animals were housed under standard vivarium conditions (22±1*°*C, 12h light/dark cycle, with ad libitum food and water). For cranial window implantation, anesthesia was first induced by 5% Isoflurane, and then maintained on 1.5-2% mixed in a gas mixture of oxygen and nitrous (analgesic) throughout the surgical procedures. When anesthetized, core body temperature of animals was maintained at 37°C. Analgesic where also administered during and post-operation.

### F. Surgical procedure

Prior to the surgeries we curve a 7 mm round glass using a similar method developed by Kim and coworkers (11). To minimize pain during surgery, inflammation and brain edema, we injected carprofen (5–10 mg/kg) and dexamethasone (2 mg/kg) subcutaneously prior to any surgical procedure. We applied ointment to the mouse’s eyes to maintain their moisture. First, the skin atop the cranium was removed and the skull surface was cleaned with a scalpel while continuously perfusing ACSF solution over the surgical area. Next, a craniotomy matched in size and shape to the outline of the 7mm glass implant was performed as follows: i) creation of a groove along the perimeter of the 7 mm round glass using a 0.5 mm diameter drill, ii) removal of the round shaped bone piece from the surrounding skull and iii) placement of the curved glass on the exposed brain, carefully avoiding touching the dura, iv) fixation of the curved glass window with super-glue to attach its edges to the surrounding skull. As final step, the window was stabilized with a thin layer of dental cement on the surrounding skull. During the drilling and exposure of the brain, artificial cerebral spinal fluid (ACSF) was constantly perfused. In order to allow head fixation during optical brain imaging, we installed a custom made annular head plate (10 mm inner diameter) around the glass window and filled the gap between the head plate and the skull with dental cement. We transferred the mouse to a recovery cage and placed it on a heating pad until it awoke. We then returned the mouse to its home cage. During the first three days following surgery, mice were subcutaneously administered with carprofen (5–10 mg/kg). Mice were allowed to recover for a period of three weeks before imaging. All data presented here was obtained from awake, head-restrained animals that were allowed to run on a carousel during imaging sessions. Prior to imaging, animals were habituated to the imaging conditions across five consecutive days.

### G. Fly preparation

GH146-Gal4 driver line was used to drive expression of GCaMP6f for volumetric calcium imaging, or ASAP2f for planar voltage imaging. Cuticle and trachea were removed, and the exposed brain was superfused with carbogenated solution (95% *O*_2_, 5% *CO*_2_) containing 103 mM *N aCl*, 3 mM *KCl*, 5 mM trehalose, 10 mM glucose, 26 mM *NaHCO*_3_, 1 mM *NaH*_2_*P O*_4_, 3 mM *CaCl*_2_, 4 mM *MgCl*_2_, 5 mM N-Tris (TES), pH 7.3. Odors at 10*-*1dilution were delivered by switching mass-flow controlled carrier and stimulus streams (CMOSense Performance Line, Sensirion) via software controlled solenoid valves (The Lee Company) or by a custom air puffer based on 2V025-08 solenoid valves. Flow rates at the exit port of the odor tube were 0.8*l/min*.

### H. Planar intravital calcium imaging

The two-photon laser scanning microscopy setup is based on a custom-modified movable objective microscope (MOM, Sutter Instrument Company) sourced by an 80 MHz 140 fs Ti:Sapphire laser (Chameleon Ultra II, Coherent Inc.) tuned to 940 nm. Laser beam power was adjusted using an electro-optic modulator (D7v-T3, Qubig GmbH) that was mounted on a multi-axis tilt platform (850-0010, EKSMA Optics) in between two polarizing beam splitters (PBS102, Thorlabs Inc.), and fed by a high voltage amplifier (A-304 customized to 200x gain, A.A. Lab Systems Ltd.). Since the scanning software (ScanImage, Vidrio Technologies, LLC.) outputs strictly positive voltages up to 2 Volts, an offset voltage of −200 Volts was added by the high voltage amplifier, to maintain a balanced dynamic range of *±*200 Volts. An achromatic quarter wave plate (AQWP10M-980, Thorlabs Inc.) introduced a bias retardation so as to minimize the optical transmission for an input voltage of −200 Volts.

An 8 kHz resonant-galvo scanning system (RESSCAN-MOM, Sutter Instrument Company) was used for planar beam steering. A 10 *×;*, 0.6NA objective lens (XLPLN10XSVMP, Olympus Corporation) was used both for excitation and collection. The collected light was directed at a fast GaAsP PMT (H10770PA-40SEL, Hamamatsu Photonics K.K.) through two dichroic mirrors (BrightLine FF735-Di01-25×36, Semrock and 565dcxr, Chroma Technology Corporation) and a bandpass filter (525/70-2P, Chroma Technology Corporation).

For PySight acquisition, the output cable of the PMT was connected directly to a high-bandwidth preamplifier (TA1000B-100-50, Fast ComTec GmbH). The amplified signal was then conveyed to a fast analog input of an ultra-fast multiscaler (MCS6A-2T8, Fast ComTec GmbH). Another analog input channel was dedicated for attenuated TTL pulses from the scanning software (ScanImage, Vidrio Technologies, LLC.), which were configured to be logically high when the galvanometric mirror scanned through the FOV. An optional frequency divider with an on-board comparator (PRL-260BNT-220, Pulse Research Lab) was used to convert the readings of the Ti:Sapphire laser’s internal photodiode into a 10 MHz clock signal, fed into the multiscaler’s reference clock input.

For analog acquisition, the output cable of the PMT was connected directly to a high-speed current amplifier (DHCPA-100, FEMTO Messtechnik GmbH). The preamplifier was DC-coupled, set to a bandwidth of 80 MHz, on its high gain setting and the output’s full bandwidth was used. The amplified signal was then conveyed to a National Instruments’ FlexRIO (PXIe-1073) digitizer with the NI 5734 adapter module, set to 120 MHz sampling frequency.

During the imaging experiment, a 440 *×;* 440*µm*_2_ FOV was imaged at 15.24 Hz 200 *µm* below the pia for 60 seconds. The PMT’s gain was adjusted to produce the highest available SNR for its respective acquisition scheme (analog or PySight), calculated in real time. Immediately afterwards the acquisition scheme was altered by rerouting the PMT’s output. Once connected the same FOV was re-imaged for the same period of time using the same parameters. The only difference between the two schemes was the PMT’s gain, which is calibrated to a higher value when imaging using the multiscaler due to its built-in discriminators. This allows the experimenter to filter out much of the multiplicative PMT noise while retaining high photosensitivity.

### I. Volumetric intravital calcium imaging

For volumetric imaging, a tunable acoustic gradient index of refraction lens (TAG Lens 2.5, TAG Optics Inc.) was inserted into the beam path upstream of the resonant-galvo system. An iris diaphragm (SM1D12CZ, Thorlabs Inc.) was used to reduce the beam diameter below 4 mm at the entrance to the TAG lens, in accordance with the effective clear aperture of the TAG lens at its 189 kHz resonance. A pair of achromatic doublet lenses with focal lengths of 45 and 60 mm (AC254-045-B and AC254-060-B, Thorlabs Inc.) were used as a relay system between the TAG lens and the resonant-galvo system. The TAG lens, its upstream iris and downstream relay system were mounted on top of a pair of manual linear translation stages (M-423, Newport Corporation) and a high load lab jack (281, Newport Corporation) for easy alignment. The TAG lens driver (DrvKit 3.3, TAG Optics Inc.) was configured to output 44 ns long TTL synchronization pulses once per focal oscillation. These were attenuated through two 20 dB RF attenuators (27-9300-20 Cinch Connectivity Solutions), before being conveyed to a fast analog input of the ultrafast multiscaler.

Before each volumetric imaging session, the laser beam was first aligned with the microscope with the TAG lens turned off and its upstream iris diaphragm fully open, using a dilute solution of fluorescein in double distilled water as an alignment target. The aperture of the iris diaphragm was then reduced to 4 mm and the TAG lens driven to the maximally permitted amplitude for its 189 kHz resonant frequency (62% of its maximal driving voltage, corresponding to an optical power difference of 20 diopters). A fixated sample of 1000 nm fluorescent microspheres, prepared according to (4), was then imaged at high digital magnification for fine alignment of the TAG lens with the beam path. As long as the lens is misaligned with the beam, the image of the beads seems to drift laterally as the objective lens is translated up and down. The micrometers of the lens’s translation stages were thus carefully driven until the said aberrations were minimized. Once aligned, the volumetric microscope was used to acquire a reference volumetric movie of a dilute solution of fluorescein in double distilled water, collapsed over time. Each acquired volumetric *Drosophila* movie could then be divided by the reference volume, to arrive at fairly accurate normalized brightness levels for each voxel. This normalization step is necessary due to the sinusoidal profile of the TAG lens focal oscillation, giving rise to uneven illumination conditions. In practice, given the 3*×;* digital zoom used for *Drosophila* volumetric imaging, the reference volume could be collapsed to a one-dimensional axial intensity variation by which the acquired 4D volumetric *Drosophila* movie was divided. Notably, for scattering tissues such as living mammalian brain, the reference volume of fluorescein solution should be mixed with scattering elements so as to arrive at comparable turbidity.

An effective axial range of 330*µm* was consistently measured with our volumetric microscope when driving the TAG lens at the said maximally permitted amplitude for its 189 kHz resonant frequency (62% of its maximal driving voltage, corresponding to an optical power difference of 20 diopters). This axial range was measured first by acquiring two z-stacks of a fixated sample of 1000 nm fluorescent microspheres, with the TAG lens on and off. It was later confirmed by manually registering features resolved in PySight-generated volumes to the respective features in a conventionally acquired z-stack. Specifically, during the imaging session on which Figures 3 and S3 are based, conventional z-stacks of 1000 nm microspheres and of two other fruit flies were acquired, along with PySight-generated volumes of the same fields of view, thereby validating that an axial range of 330*µm* was available in this particular imaging session as well.

Adult female *Drosophila melanogaster* were prepared as detailed above. Each trial lasted 33.5 seconds (2462 volumes at 73.4 volumes per second). After five seconds of baseline imaging, a five-seconds long puff of 2-Pentanone or isoamyl acetate was delivered to the fly. After five additional seconds, an additional, identical, five-seconds long puff of the same odor was delivered to the fly. A proximal negative pressure air inlet continuously cleared the chamber from odor traces throughout the imaging session. Six trials were acquired for each odor, with an inter-trial interval of 2-4 minutes. The imaged FOV was shaped as a 200:512 rectangle at a digital magnification of 3*×;*, which corresponds to 234 *×;*600 *×;*330*µm*_3_ using our 10*×;* objective lens and TAG lens. The resulting imaging rate was 73.4 volumes per second. We then repeated the experiment while zooming in on the antennal lobes, using a FOV shaped as a 220:512 rectangle at a digital magnification of 7*×;*, which corresponds to 110 *×;*257 *×;* 330*µm*_3_ using our 10 *×;* objective lens and TAG lens. The resulting imaging rate was 67.2 volumes per second.

### J. Planar intravital voltage imaging

Four adult female *Drosophila melanogaster* were prepared as detailed above. The lateral horn, mushroom body and antennal lobe were imaged by TPLSM using a Sutter Instrument Company’s DF scope with resonant scanners (8 kHz, RESSCAN-MOM) based on an Olympus BX51WI microscope, Mai Tai HP DS laser. The imaging software used in the DF-scope was Sutter’s MScan. Before imaging, the line signal from the resonant mirrors was connected to an analog input of the multiscaler. This doesn’t affect the performance of the imaging system in any way. For each fly, between five and ten 30-seconds-long imaging sessions were conducted using the same acquisition scheme (analog or photon-counting). The size of the FOV was 10 *×;* 50*µm* to allow for imaging rates of 247 Hz. After these initial repetitions, the output of the PMT would be rerouted either to a fast amplifier (TA1000B-100-50, Fast ComTec) connected to the multiscaler (to switch the scheme to a photon-counting one) or to a slower preamplifier (C7319, Hamamatsu) connected to a National Instruments’ FlexRIO digitizer, and the experiment could be re-executed with the same overall parameters. Data acquired using both the multiscaler and MScan was converted to a .tif format and processed using identical custom Python scripts available upon request. The analysis first required the user to mark the region of interest (ROI) containing the anatomical structure in question. The output consisted of the mean fluorescence trace inside the ROI across all five to ten repetitions of each fly, as well as the individual fluorescence traces per repetition per animal.

### K. Data analysis

#### Planar intravital calcium imaging

Data acquired using both acquisition schemes was converted to a standard.tif format and analyzed with the CaImAn framework (18) using the same parameters. Results were processed using custom Python scripts available on request. To compare the mean Δ*F/F* values as well as the different rise times, a two-sample, two-sided t-test with Welch’s correction was used.

#### Volumetric intravital calcium imaging

Each trial was parsed by PySight to a 4D volume consisting of 2462 *×;* 200 *×;* 512*×;*150 voxels in *txyz*. Its intensity profile along the axial dimension was normalized by the axial intensity profile acquired in a dilute fluorescein solution as elaborated above. A cuboid volumetric region of interest was manually selected for each olfactory structure, and the normalized brightness of all of its voxels were evenly summed. The median brightness in the first 4.7 seconds (346 volumes) of each trial was considered to be the baseline brightness. Fluorescence variations were then calculated by subtracting the baseline brightness from the instantaneous brightness, and dividing the result by the baseline brightness.

The size of the list files for the twelve trials ranged between 161,851,819-164,939,179 and 169,968,217-170,897,305 bytes for isoamyl acetate and 2-Pentanone trials, respectively. The mean data throughput was calculated by dividing the size of the largest list file (170,897,305 bytes) by the total acquisition length (33.54 seconds), neglecting list file header length.

Similarly for volumetric imaging of the antennal lobes, each trial was parsed by PySight to a 4D volume consisting of 2254 *×;* 220 *×;* 512 *×;* 150 voxels in *txyz*. Its intensity profile along the axial dimension was normalized by the axial intensity profile acquired in a dilute fluorescein solution as elaborated above. The median brightness in the first 5 seconds (335 volumes) of each trial was considered to be the baseline brightness. Fluorescence variations were then calculated by subtracting the baseline brightness from the instantaneous brightness, and dividing the result by the baseline brightness. An ellipsoid volumetric region of interest was manually selected for two glomeruli (A and B), and the normalized brightness of all of their voxels were evenly summed. Conversely, to identify glomeruli C that were more responsive to isoamyl acetate than to 2-Pentanone, we first binned the 4D volume from each of the 12 trials 2×2×;2 times in *xyz* to reduce the computational load. We then summed the intensity-corrected brightness along 5.5 seconds following the first odor puff onset, and applied a 3D Gaussian filter with a standard deviation of 3×3×;1 voxels (3*×;* 3*×;* 4.4*µm*_3_) for each individual trial. Voxels within the antennal lobes were considered to have a preferential response to isoamyl acetate if their minimal brightness across isoamyl acetate odor puff trials was higher than the maximal brightness across 2-Pentanone odor puff trials. The binary mask of odor-preferential voxels was applied on the trial-summed (unsmoothed, full-sized) 4D volume. Its time-collapsed 3D volume was rendered in AMIRA 6.4 (Thermo Fisher Scientific) yielding Fig 3d, and its respective temporal fluorescence variation traces (Figure 3e-f) were calculated using Python scripts available upon request. A sequential montage of antennal lobe slices in different axial planes was prepared by binning the trial-summed 4D volume 336×4×;4×3 times in *txyz*, then displaying every sixth slice along the axial dimension, resulting in 6.6 *µm* thick axial slices evenly spaced 39.6 *µm* apart. Color intensity was normalized for each color mask separately, thereby artificially highlighting the glomeruli of interest.

### L. Software

PySight is an open-source multi-dimensional photon counting rendering engine written in Python, which can be installed via the Python pip application: `pip install pysight`. The full source code can be found at https://github.com/PBLab/python-pysight, published under the Creative Commons license.

## Acknowledgments

PB and MP both thank the support from ERC projects 639416 and 676844 respectively. PB also thanks the support of the BSF foundation (grant 2014509). LG and PB thank Jonathan Driscoll for helpful discussions through the hardware selection process.

---

**Supplementary Figure S1.**
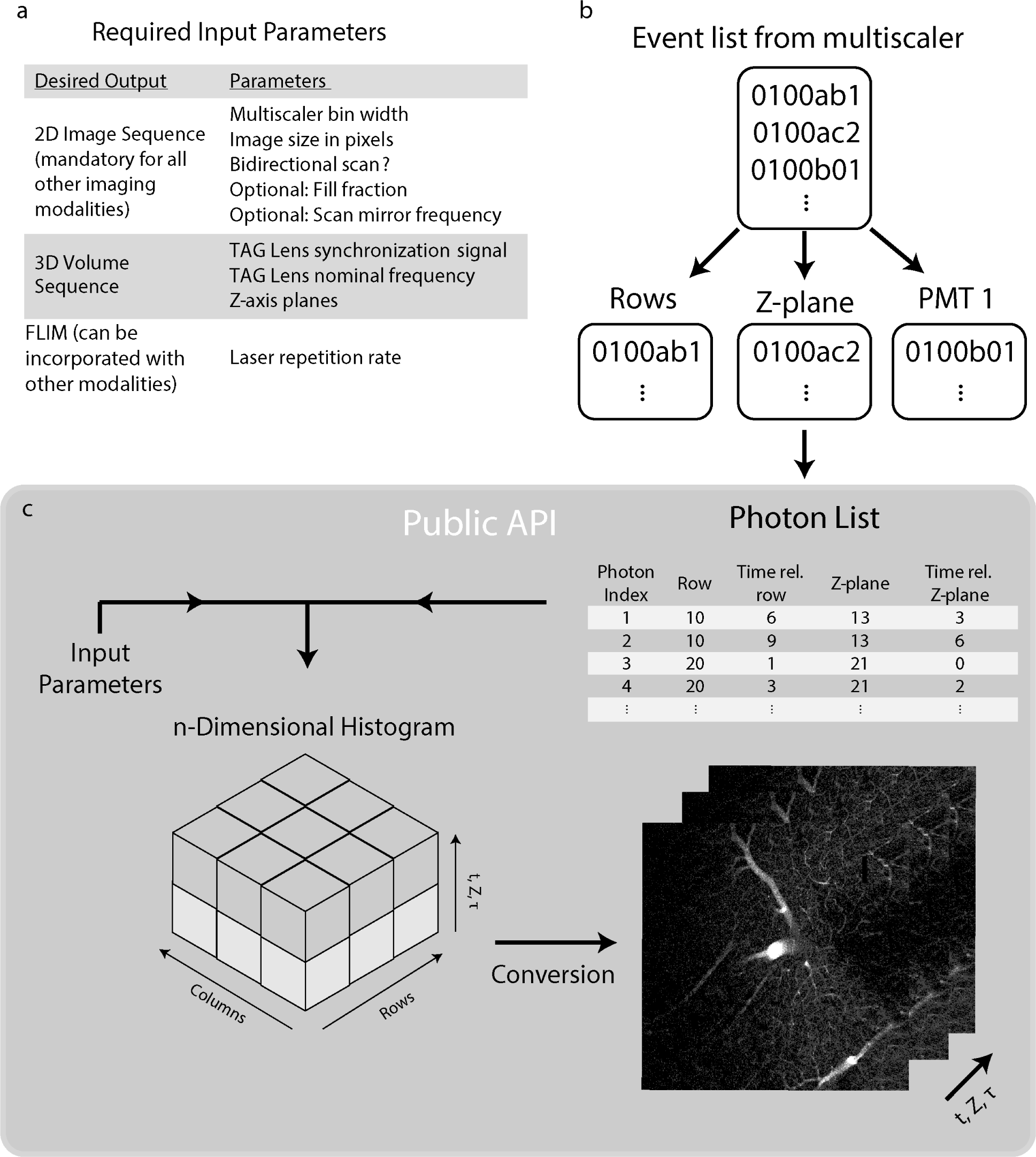
Flowchart of PySight’s logic. a) Input parameters for PySight. A small set of input parameters and synchronization signals is sufficient to generate multi-dimensional image stacks. b-c) The list of events generated by the multiscaler is demultiplexed back to its originating channels (`STOP1`, `STOP2` and `START` in Figure 1a) and converted to time values. Each event representing a photon detection is assigned to its respective coordinates along each recorded dimension. For example, in a given experiment a photon could have corresponding start-of-frame, start-of-line, depth in the sample (*z*), time-since-laser-pulse (for lifetime imaging) and a spectral channel, effectively producing a time-lapse stack of 5D images if all synchronization signal parameters are provided. The photons are binned into an n-dimensional array, such as *xyztτ C*, with *τ* and *C* corresponding to fluorescence lifetime and spectral channel, respectively. PySight can also generate a multi-dimensional stack of images from lists of photons not generated by a MCS6A multiscaler, provided that they comply with the input format.

**Supplementary Figure S2.**
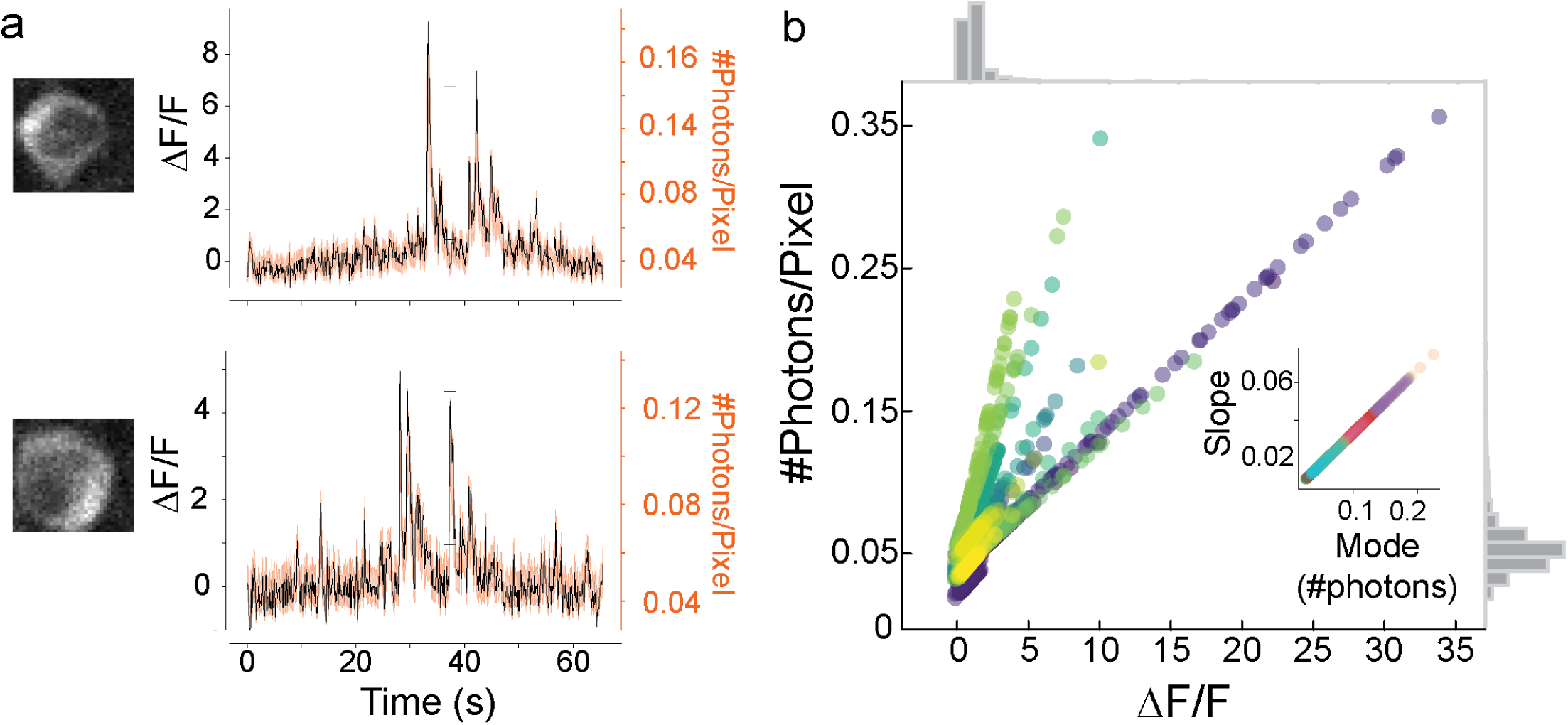
Comparison of photons counts and Δ*F/F* signals. a) Close-up view of two individual neuronal somata acquired using photon counting, with their Δ*F/F* traces (black) and a confidence interval for the number of photons the Δ*F/F* value represents (orange). An interval is required due to the non-constant amount of laser pulses arriving at each pixel. b) Summary of the expected values of Δ*F/F* in photons for all cells. As the inset suggests, each cell has a unique ratio, possible due to factors such as GCaMP concentration and slightly different collection efficiency. The data was acquired with a pixel dwell time of 44 ns.

**Supplementary Figure S3.**
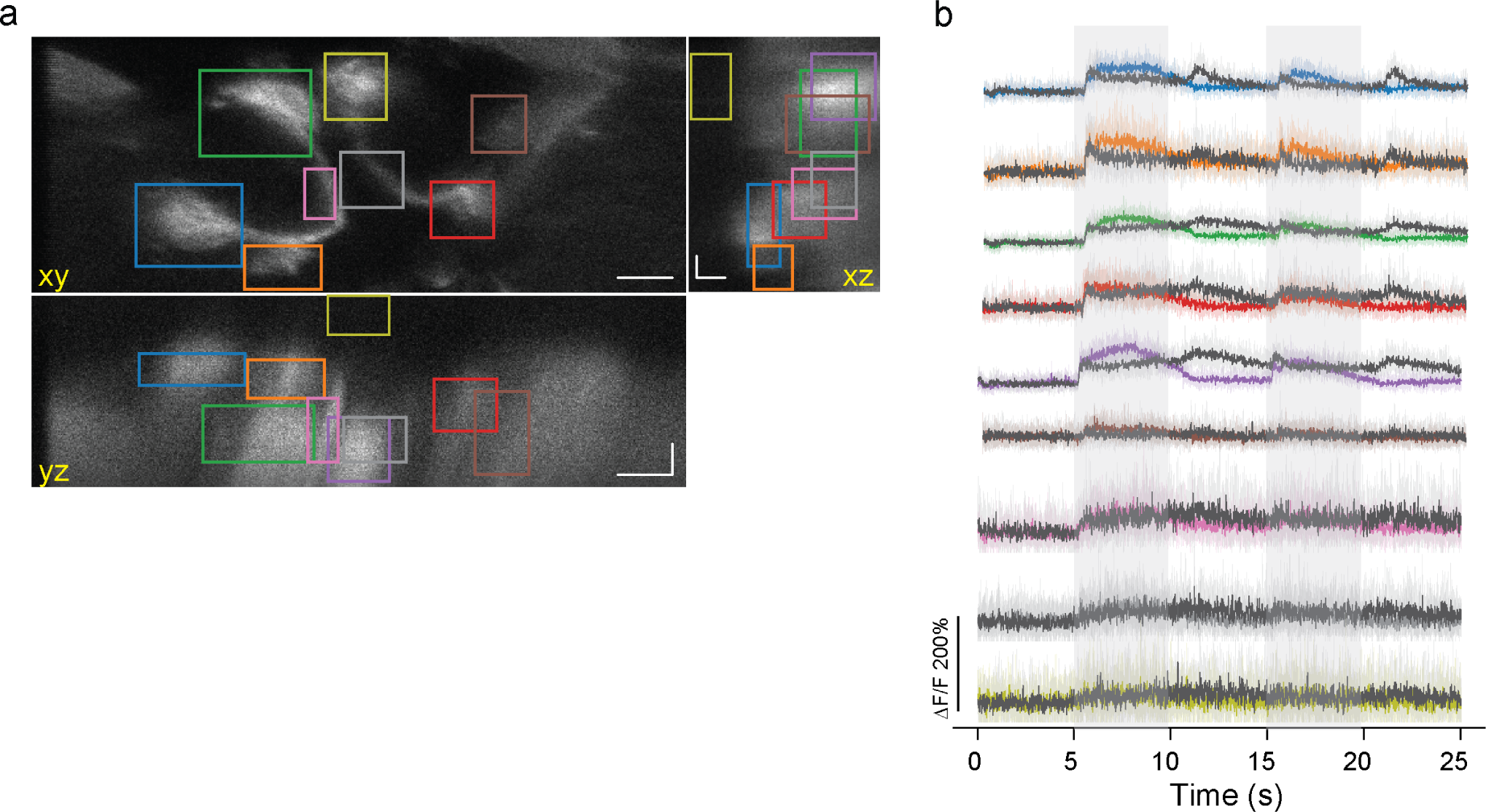
Volumetric GCaMP6f traces (Δ*F/F*) from distinct volumes of interest (VOI) in a drosophila’s olfactory brain regions were acquired at 73.4 volumes per second over a volume of 234 *×;* 600 *×;* 330*µm*^3^. The location of each VOI can be seen on the three sum-projections (across all volumes and over time). Notice the lack of response to the different odor in a VOI located in a brain region deprived of GCaMP6f expression (yellow trace, bottom in panel b).

## Supplementary Note 1: PySight-compatible hardware

While PySight has been built around specific hardware (the MSCSA, top row), its open-source code can handle any list of photon arrival times through a well-documented application interface (Supplementary Figure S1). The following non-exhaustive table compares the key features of commercially-available time-to-digital converters that could be used with PySight software.

**Table S1.** List of PySight-compatible hardware that can record photon arrival times in time-stamping mode.

| Product | Manufacturer | Deadtime<br>[ns] | Sustained<br>rate<sup>a</sup> [Mcps] | Peak rate<br>[Mcps] | Burst rate<sup>a</sup><br>[Mcps] | FIFO<sup>a</sup><br>[counts] | Resolution<br>[ps] |
| --- | --- | --- | --- | --- | --- | --- | --- |
| MCS6A | FAST ComTec | 0 | 17 | 10000<sup>b</sup> | 100 | 256M | 100 |
| TimeHarp 260 NANO | PicoQuant | <2 | 40 |  |  | 8.4M | 250 |
| FastFLIM | ISS | 3.125 | 15 |  |  |  | 100 |
| quTAU | qutools | 5.5 | 3 |  |  |  | 81 |
| SPC-160 PCIE | Becker & Hickl | 80 | 4 |  | 12.5 | 2M | 2.5 |
| HRM-TDC | SensL | 190 | 4.5 |  | 100 |  | 66 |

<sup>a</sup> The on-board FIFO, where available, is filled as rapidly as specified by the burst rate, and drained as rapidly as specified by the sustained rate.
<sup>b</sup> A peak count rate of 10 Gigacounts per second per channel can be sustained for 6.5 microseconds.

### A. Supplementary Protocol: Volumetric TPLSM with a MCS6A multiscaler

This protocol guides the prospective user through the necessary steps towards laser scanning microscopy with a MCS6A multiscaler. For your convenience, we provide customized settings files tailored for planar and volumetric two-photon microscopy in the GitHub repository that accompanies this paper (https://github.com/PBLab/python-pysight)

1. Follow the installation procedure detailed at section 2 of the multiscaler’s user manual available at FAST website.
2. Conduct the first two introductory measurements described at the ‘Getting Started’ subsection of the multiscaler. The more advanced examples available in that subsection are not essential for ordinary two-photon imaging.
3. Open the downloaded settings files with a text editor and replace the number 535 at their first line with the unit number of your particular MCS6A multiscaler.
4. Copy the provided settings files (https://github.com/PBLab/python-pysight) to the MPANT’s folder, e.g. ‘C:\MCS6A (x64)’.
5. Use the ‘MCS6A Settings’ dialogue box to load settings file ‘pre_test_2d_a.set’ (for planar imaging) or ‘pre_test_3d_a.set’ (for volumetric imaging).
6. Use the ‘Data Operations’ dialogue box to select the data taking folder and define an informative data file name, e.g. ‘D:\Data\User\Preliminary_test.mpa’. Make sure that the ‘Write Listfile’ checkbox is checked.
7. Configure your microscopy scanning software to output a pulsatile signal once per line. Use RF attenuators to attenuate the line synchronization signal to acceptable voltages, as specified in subsection 8.2 of the multiscaler’s user manual. Preferably use low absolute voltages, in the 20-50 mV range, to minimize the risk for contamination of the PMT signal. ! Critical Excessive voltages or currents may permanently damage the multiscaler, or cause suboptimal performance. Consult subsection 8.2 of the multiscaler’s user manual for the acceptable voltage range of the analog (START and STOP) inputs. Measure your signals with an oscilloscope before connecting them to the multiscaler.
8. Connect the line synchronization signal to the analog ‘START’ channel of the multiscaler.
9. Click the ‘Inputs…’ button at the ‘MCS6A Settings’ dialogue box, in order to open the ‘Input Thresholds’ dialogue box. Tune the threshold level for the line signal to about half of the amplitude of the attenuated line synchronization signal. For example, if the attenuated line synchronization signal rises to +30 milliVolts when a line scan begins, select a discriminator threshold of +15 milliVolts and a rising edge detection.
10. Start scanning with the laser shutter turned off. Start multiscaler acquisition by clicking the ► icon, or the ‘Start’ button under the ‘Action’ menu. The periodicity and total number of acquired line signals in the resulting spectrum should match the line rate of your scanning system. Otherwise, reconfigure the threshold level for the line signal.
11. Turn on your PMT in light-tight conditions, and increase its gain to near-maximal values (e.g. a control voltage of 850mV for Hamamatsu H10770PA-40). Turn on the preamplifier and measure the amplified signal with an oscilloscope. Pay attention to the polarity of your amplified signal, which varies among preamplifier models. Take note of baseline noise levels and maximal pulse height. ! Critical Excessive voltages or currents may permanently damage the multiscaler, or cause suboptimal performance. Consult subsection 8.2 of the multiscaler’s user manual for the acceptable voltage range of the analog (START and STOP) inputs.
12. Connect your pre-amplified PMT signal to the analog ‘STOP1’ channel of the multiscaler.
13. Click the ‘Inputs…’ button at the ‘MCS6A Settings’ dialogue box, in order to open the ‘Input Thresholds’ dialogue box. Tune the threshold level for the PMT signal above the baseline noise level measured earlier. For instance, if the amplified noise level through a non-inverting preamplifier is about *-120±* 15 milliVolts, a discriminator threshold of *-180* mV and a falling edge detection should capture most photodetection events while discarding most noise fluctuations. ! Critical The absolute voltage difference between any analog input signal and its selected threshold level should never exceed 2 Volts. Verify that the input thresholds you select match the polarity of your pre-amplified signals.
14. Start scanning some fluorescent solution and then start multiscaler acquisition by clicking the ► icon, or the ‘Start’ button under the ‘Action’ menu. The periodicity of the acquired photon detection events should match the line periodicity of your laser scanning system. Mark down the total number of counts throughout the measurement.
15. Turn off the laser shutter and PMT and block light from reaching your PMT. Start another multiscaler acquisition with the preamplifier turned on. If the threshold level was selected properly, no counts should be acquired.
16. Turn on the PMT in light-tight conditions and start another multiscaler acquisition. If the threshold level was selected properly, the acquired count rate should be comparable to the typical dark count rate of your PMT. Otherwise, reconfigure your threshold level and repeat steps 14-16. ! Critical A sufficiently high threshold level leads to the rejection of most dark counts, at the price of rejecting a considerable percentage of legitimate photon detection events. The total number of counts throughout the measurement (as measured in step 14) should only weakly depend on the threshold level, if selected properly. Particularly, if the total number of counts varies by more than 8% following a 10% change in the threshold value, then the threshold value is probably wrong.
17. Repeat steps 11-16 for any additional PMT, connecting it to additional analog ‘STOP’ channels of the multiscaler.
18. Optional: For volumetric imaging, take the following additional steps:

a. Configure your TAG lens driver to output a pulsatile signal once per lens oscillation, preferably at 0*°* or 180*°* phase.
b. Use RF attenuators to attenuate the TAG synchronization signal to acceptable voltages, as specified in subsection 8.2 of the multiscaler’s user manual. Preferably use low absolute voltages, in the 20-50 mV range, to minimize the risk for contamination of the PMT signal. ! Critical Excessive voltages or currents may permanently damage the multiscaler, or cause suboptimal performance. Consult subsection 8.2 of the multiscaler’s user manual for the acceptable voltage range of the analog (START and STOP) inputs. Measure your signals with an oscilloscope before connecting them to the multiscaler.
c. Connect the TAG synchronization signal to the analog ‘STOP2’ channel of the multiscaler.
d. Click the ‘Inputs…’ button at the ‘MCS6A Settings’ dialogue box, in order to open the ‘Input Thresholds’ dialogue box. Tune the threshold level of the TAG synchronization signal to about half of the amplitude of the attenuated TAG synchronization signal. For example, if the attenuated TAG synchronization signal rises to +30 milliVolts when a TAG lens oscillation begins, select a discriminator threshold of +15 milliVolts and a rising edge detection.
e. Lock the TAG lens at the desired resonant frequency, with the laser shutter turned off. Once locked, start multiscaler acquisition by clicking the ► icon, or the ‘Start’ button under the ‘Action’ menu. The periodicity of acquired TAG synchronization signals in the resulting spectrum should match the periodicity of the TAG lens. Otherwise, reconfigure the threshold level for the TAG synchronization signal.
19. Open the ‘MCS6A Settings’ dialogue box and increase the bin width as necessary to obtain a sufficiently long acquisition for your experiment. The bin width and resulting histograms do not influence offline processing by PySight. You may thus check the ‘No Histogram’ box to reduce computational overhead during automated imaging sessions. Online monitoring of the acquired histogram is helpful however for fault detection and troubleshooting.
20. The multiscaler is now optimized for data acquisition with your particular hardware. Use the ‘MCS6A Settings’ dialogue box to save settings file ‘My_system_settings_A.SET’, which you could load from now on before each imaging session.
21. Optional: For online monitoring of the acquired image through an existing data acquisition card, connect the fast ‘SYNC 1’ output from the front panel of the multiscaler to an input port of your data acquisition hardware. Use the ‘Sync1’ drop-down menu within the settings dialogue box to select which analog input channel to broadcast through the ‘SYNC 1’ output channel. Image as usual.

